# Integration of Contours Defined by Second-Order Contrast-Modulation of Texture

**DOI:** 10.1101/792820

**Authors:** Alex S Baldwin, Madeleine Kenwood, Robert F Hess

## Abstract

Boundaries in the visual world can be defined by changes in luminance and texture in the input image. A “contour integration” process joins together local changes into percepts of lines or edges. A previous study tested the integration of contours defined by second-order contrast-modulation. Their contours were placed in a background of random wavelets. Subjects performed near chance. We re-visited second-order contour integration with a different task. Subjects distinguished contours with “good continuation” from distractors. We measured thresholds in different amounts of external orientation or position noise. This gave two noise-masking functions. We also measured thresholds for contours with a baseline curvature to assess performance with more curvy targets. Our subjects were able to discriminate the good continuation of second-order contours. Thresholds were higher than for first-order contours. In our modelling, we found this was due to multiple factors. There was a doubling of equivalent internal noise between first- and second-order contour integration. There was also a reduction in efficiency. The efficiency difference was only significant in our orientation noise condition. For both first- and second-order stimuli, subjects were also able to perform our task with more curved contours. We conclude that humans can integrate second-order contours, even when they are curved. There is however reduced performance compared to first-order contours. We find both an impaired input to the integrating mechanism, and reduced efficiency seem responsible. Second-order contour integration may be more affected by the noise background used in the previous study. Difficulty segregating that background may explain their result.

## 1 Introduction

The human visual system extracts information from its input (the retinal images from the two eyes) to drive behaviour. In that input, the “difference[s] that make a difference” (Bateson, 1970) occur at the boundaries of objects, surfaces, and textures (Attneave, 1954; Elder, 1999). Many of these boundaries result in a change in luminance (for example, the edge of a light object on a dark background). These can be encoded by classical “edge-detecting” mechanisms (Marr, 1976). This is thought to be one role of the simple cells found in primary visual cortex. They have receptive fields tuned to detect local luminance changes. The changes they detect would correspond to oriented bars or edges in the input image (Hubel and Wiesel, 1959). The features defined by local changes in luminance (i.e. boundaries between light and dark) are classified as “first-order” features. Take for example a first-order sinusoidal grating. Its luminance varies across space. The amplitude of the variation determines the grating’s contrast. The variation is sinusoidal, with a particular spatial frequency, along the orientation of its axis. When the Fourier transform is taken of that grating, it is represented as a point in the output. The position of that point is determined by the grating’s spatial frequency and orientation. It has been suggested that visual processing involves a decomposition of the input into its Fourier components (Graham, 1981). Under this proposal, such first-order features are readily detected. Beyond these however, there are then a range of “second-order” features. These are defined by differences in the local first-order properties (Cavanagh and Mather, 1989). For example, a second-order boundary can occur between regions of higher and lower texture contrast (Schofield, 2000; Johnson and Baker, 2004). Second-order features require additional processing to be represented clearly in the Fourier transform (Henning et al., 1975; Chubb and Sperling, 1988). This additional processing must be non-linear. A suitable non-linear pathway in the visual system has been identified with electrophysiology (Zhou and Baker, 1993, 1994; Mareschal and Baker, 1998). Results from the macaque (Li et al., 2014) show a population of neurones that multiplex first- and second-order responses (with similar receptive field tunings). Other evidence comes from behavioural studies. Psychophysical experiments have explored the relationships between processing channels for stimuli defined by luminance-modulation (first-order) and contrast-modulation (second-order) (e.g. Sutter et al., 1995; Schofield and Georgeson, 1999). The results from these experiments have been used to formulate models of second-order processing. These typically sandwich a nonlinear stage (e.g. rectification) between two filtering stages (further examples in Graham et al., 1992, 1993; Graham and Sutter, 1998; Landy and Oruç, 2002).

There is a retinotopic mapping of the visual input to the early visual cortical areas. The neurones in these areas directly encode the local properties of the input. For example, the orientation and position of individual parts of a continuous contour. Each neurone can only represent the presence of a small part of the contour. The representation of the whole requires a combination of responses across multiple neurones. Behaviourally, we can investigate the psychophysical analogue of this “contour integration” process. A landmark study by Field et al. (1993) introduced a paradigm to do so. This approach has since been adapted to look at a wide range of questions regarding the perception of contours. In the Field et al. (1993) paradigm, a target contour was presented composed of a series of discrete wavelets (similar to that shown in Figure 1A). These wavelets are presented in a background “texture” of randomly-oriented wavelets. Subjects were tested on their ability to detect the target contour. The task had them discriminate stimuli containing a target from those containing only the background texture. The original study put forward a theory for how contours are integrated. They suggested that adjacent wavelets in the contour are joined-up based on an “association field” (Field et al., 1993). This defines the set of wavelets that can be linked together. Whether a wavelet can be linked is determined by its combined orientation and position information. Sophisticated models of contour integration have been developed based on ecological parameters (Geisler et al., 2001), psychophysical results (Hess et al., 1998; Ledgeway et al., 2005; May and Hess, 2007, 2008), and knowledge of underlying physiology (Li, 1998; Angelucci et al., 2002; Field and Hayes, 2004; Li et al., 2006). The body of work in this area has recently been reviewed by Hess et al. (2015).

**Figure 1:**
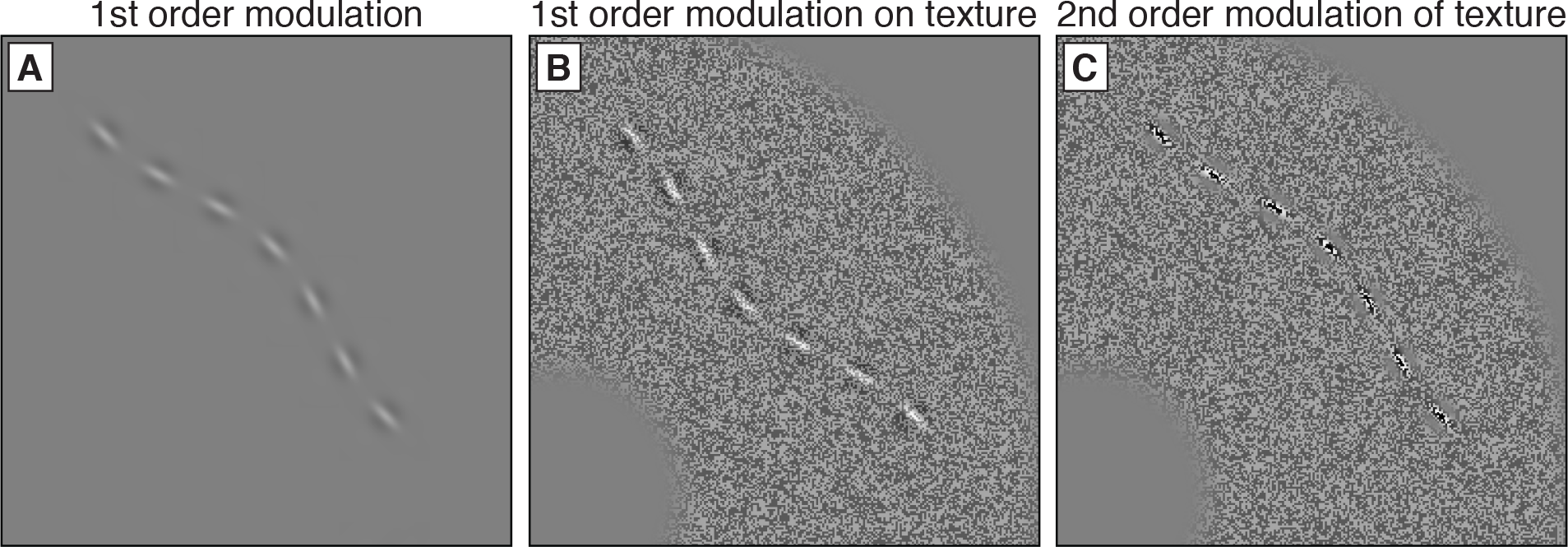
Examples of the three types of contours used in this study. Panel A shows the 1st order contour (at 30% contrast), similar to that used in the previous study with this method (Baldwin et al., 2017). Panel B shows a 1st order contour (65% contrast) added on top of the binary noise texture (30% contrast) used in the experiments . Panel C shows a contour formed of 2nd order wavelets that modulate the contrast of that noise (260% modulation). The curvature modulation amplitude of each contour was 2 cycles.

If presented without the random wavelet background, the contour stimuli used in the Field et al. (1993) task would be trivial to detect. The task becomes difficult when the contours are displayed in the random wavelet background. This background contains spurious contours that interfere with performance in the task (Watt et al., 2008). There is a clear ecological validity for this task, which tests the ability to segregate a contour from a noisy background. More recently however, we have been interested in the basic behaviour of the process that links the local responses (by their “good continuation”). For this reason, we have recently developed a new task (Baldwin et al., 2017). In our new task we measure the ability of subjects to discriminate the good continuation of contours. We do not rely on adding random noise to the stimuli to degrade performance. Instead, the target contour must be discriminated from distractor contours. Like Field et al. (1993), our contours are made up of discrete wavelets. The target contour wavelets have positions and orientations consistent with an underlying (imaginary) curve. This curve can flex in either direction. The positions and orientations of the wavelets in the distractor contours are drawn from the same sets as those used in the targets. The distinction is in the relationship between position and orientation. The wavelets in the distractor contour have orientations consistent with the *opposite* direction of curvature. Subjects must therefore combine position and orientation information from the wavelets to discriminate the target contour.

The task presented in Baldwin et al. (2017) does not rely on background noise to measure thresholds. It is theoretically possible for an ideal observer to discriminate the target contour perfectly. This allows noise to be introduced back into the task to see how it affects discrimination thresholds. Thresholds can be measured as a function of external noise standard deviation. This gives a noise-masking function. Noise-masking functions have been measured for a wide variety of visual tasks. For a contrast detection task, performance in contrast noise can be measured (Pelli, 1981). Similar approaches have been taken to the discrimination of texture orientation (Dakin, 2001), motion (Dakin et al., 2005) and stereo (Wardle et al., 2012). These noise-masking functions are typically interpreted through model-fitting. The fitted model parameters explain the pattern of thresholds as resulting from multiple factors (Pelli and Farell, 1999; Lu and Dosher, 2008). The Linear Amplifier Model (Eq. 3 in Methods section) is the simplest of these models. In the Linear Amplifier Model, thresholds are determined by a combination of two parameters. These are the equivalent internal noise (*σ*_eq.int_) and the processing efficiency (*β*). When relatively low levels of external noise are added to the input stimulus there is no effect on thresholds. At this point performance is said to still be limited by the “internal noise” in the visual system (Gregory and Cane, 1955; Barlow, 1977). At some critical level of external noise thresholds begin to increase. The equivalent internal noise is defined as the amount of external noise that must be added to reach this point. It is worth noting that differences measured in this “equivalent internal noise” can be explained by a variety of input-level factors. They are not necessarily attributable to actual changes in the amount of internal noise in the system (Baldwin et al., 2016).

The second parameter in the Linear Amplifier Model is the processing efficiency. This accounts for how well the subject is able to make use of the noisy stimulus information (Barlow, 1977). This parameter can be placed on an absolute scale by comparison to the ideal observer (Geisler, 2004). Making this comparison requires the derivation of an ideal observer prediction specific to the task. The ideal observer gives a benchmark performance for a specific external noise level. Discrepancies between human and ideal performance can have a variety of explanations. A previous study explored the discrimination of the orientation of textures formed of wavelets. In that study, human efficiency was presented as the effective “number of wavelets” whose information was used to perform the task (Dakin, 2001). This assumes that the subjects use the information from that subset of wavelets ideally. We have previously conducted a noise-masking study on contour integration (Baldwin et al., 2017). We measured thresholds as a function of three different types of external noise variance. We found that contrast noise (randomising the contrasts of the wavelets) had no effect on performance. Adding orientation noise or position noise resulted in typical noise-masking functions. From those noise-masking functions we determined the equivalent internal noise for orientation (6°) and for position (3 arcmin). We also developed an ideal observer for the task. This allowed us to report human efficiency relative to the ideal (36% in orientation noise, 21% in position noise). A subsequent experiment used that task to test patients with mild traumatic brain injury. Those patients showed elevated equivalent internal noise (Ruiz et al., 2019).

In this study we are interested in the integration of contours composed of second-order wavelets (e.g. Figure 1C). These wavelets are created through second-order modulation of texture contrast. A previous study using the Field et al. (1993) method found that contours formed from second-order wavelets were not integrated (Hess et al., 2000). Subjects’ performance for discriminating contours in that study was close to chance. Where subjects did perform above-chance in that study, this was attributed to local interactions between a couple of wavelets. They did not find evidence for global integration into a contour. This result is surprising when compared to results found in other tasks. Subjects *can* perceive shapes that are defined by second-order modulation (Hess et al., 2001; Bell and Badcock, 2008). Those studies used radial-frequency pattern shapes with continuous outlines. These may have an advantage compared to the Hess et al. (2000) contours, which were divided into discrete wavelets. The contour integration result can also be compared to Baldwin et al. (2015). That study tested judgements of coherent texture orientation (Husk et al., 2012). They found that such judgements could be made for stimuli defined by second-order wavelets. In fact, efficiency for those stimuli (81%) was not much lower than that for first-order stimuli (98%). A separate study developed a model to account for performance in the coherent texture orientation task (Baldwin et al., 2014). This model had computations performed using mechanisms with relatively large receptive fields. It is possible that this task may make use of second-order mechanisms that cannot be taken as inputs for contour integration. Here, we take another look at the integration of contours composed of second-order contrast-modulation. We use our recently-developed psychophysical task with a noise-masking paradigm. This allows us to compare the equivalent internal noise and processing efficiency parameters for first- and second-order contours. We also extended our task to investigate the integration of more curvy contours (to examine whether second-order contour integration breaks down under these conditions). Finally, we compare our results to those obtained from an orientation discrimination experiment.

## 2 Methods

### 2.1 Apparatus

The experiment was programmed in Matlab (version R2016b) and presented using Psychtoolbox (Brainard, 1997). Stimuli were displayed on a Dell P1130 monitor with a resolution of 1280 × 1024 pixels and a framerate of 85 Hz. A Bits# Stimulus Processor (Cambridge Research Systems Ltd., Kent, UK) allowed us to present stimuli with a 14-bit luminance resolution. The mean luminance was 82 cd/m^2^. Subjects sat at a viewing distance of 86 cm. At this distance, there were 50 pixels per degree of visual angle.

### 2.2 Participants

Four subjects took part in this study (two female, with an age range of 21-35). Subject details are given in Table 1 The subjects numbered S2 and S3 were authors. All subjects had either normal or corrected-to-normal vision. The subjects gave written informed consent. All experiments were approved by the Research Ethics Board of the McGill University Health Centre, and carried our in accordance with the Declaration of Helsinki.

**Table 1:** Information for the four subjects. This includes the thresholds from the contrast (1^st^ order and 1^st^ order on noise texture) and modulation (2^nd^ order modulation of noise texture) detection tasks that were used to set the stimuli in the contour integration experiments to be presented at three times the threshold level.

| Subject # | Age | Gender | Detection thresholds (%) [with 95% CI] |  |  |
| --- | --- | --- | --- | --- | --- |
| | | | 1<sup>st</sup> order | 1<sup>st</sup> order + texture | 2<sup>nd</sup> order × texture |
| S1 | 35 | M | 3.6; [2.5 - 4.9] | 21.2; [18.0 - 24.3] | 65.0; [52.3 - 78.4] |
| S2 | 30 | M | 2.6; [2.3 - 2.9] | 21.1; [18.6 - 24.2] | 65.7; [57.7 - 75.3] |
| S3 | 21 | F | 2.2; [2.0 - 2.5] | 18.5; [15.1 - 21.8] | 71.4; [63.2 - 78.8] |
| S4 | 21 | F | 2.4; [2.1 - 2.6] | 20.2; [17.8 - 22.6] | 64.1; [53.5 - 74.4] |

### 2.3 Stimuli

Contours were made up of seven log-Gabor wavelets (Meese, 2010; Baldwin, 2013) placed on a path defined by a cosine function (Figure 1). Detailed methods for stimulus generation are provided in our previous publication (Baldwin et al., 2017). The wavelets had a spatial frequency of 1.8 cycles per degree of visual angle. They had a spatial frequency bandwidth of 1.6 octaves, and an orientation bandwidth of ±30°. All on-screen distances are given in cycles (periods of the stimulus spatial frequency), where one cycle is 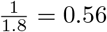 degrees of visual angle. The distance between the first and last wavelet of each contour was 14 cycles. The centre of the contour was 10.8 cycles from the fixation point in the centre of the display. The orientation of the wavelets depended on whether they linked to form a target or a non-target distractor contour. For target contours, the orientations lined up with the local path of the contour. For non-target contours, the orientations were reversed. The non-target contour orientations would be correct for a contour curving in the *opposite* direction.

In our noise-masking experiments, we also generated contours with added external noise. For the orientation noise we added a random orientation offset to every wavelet. These offsets were drawn from a normal distribution with a mean of zero and a standard deviation equal to the external noise level. For the position noise we added random horizontal and vertical shifts. These were also drawn from a normal distribution.

The wavelets forming the contour were defined by either luminance-modulation or texture contrast-modulation. First-order stimuli were modulated about the mean luminance of the display (Figure 2B-C). The contrast of the wavelets was defined as delta-contrast

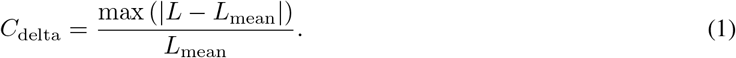

**Figure 2:**
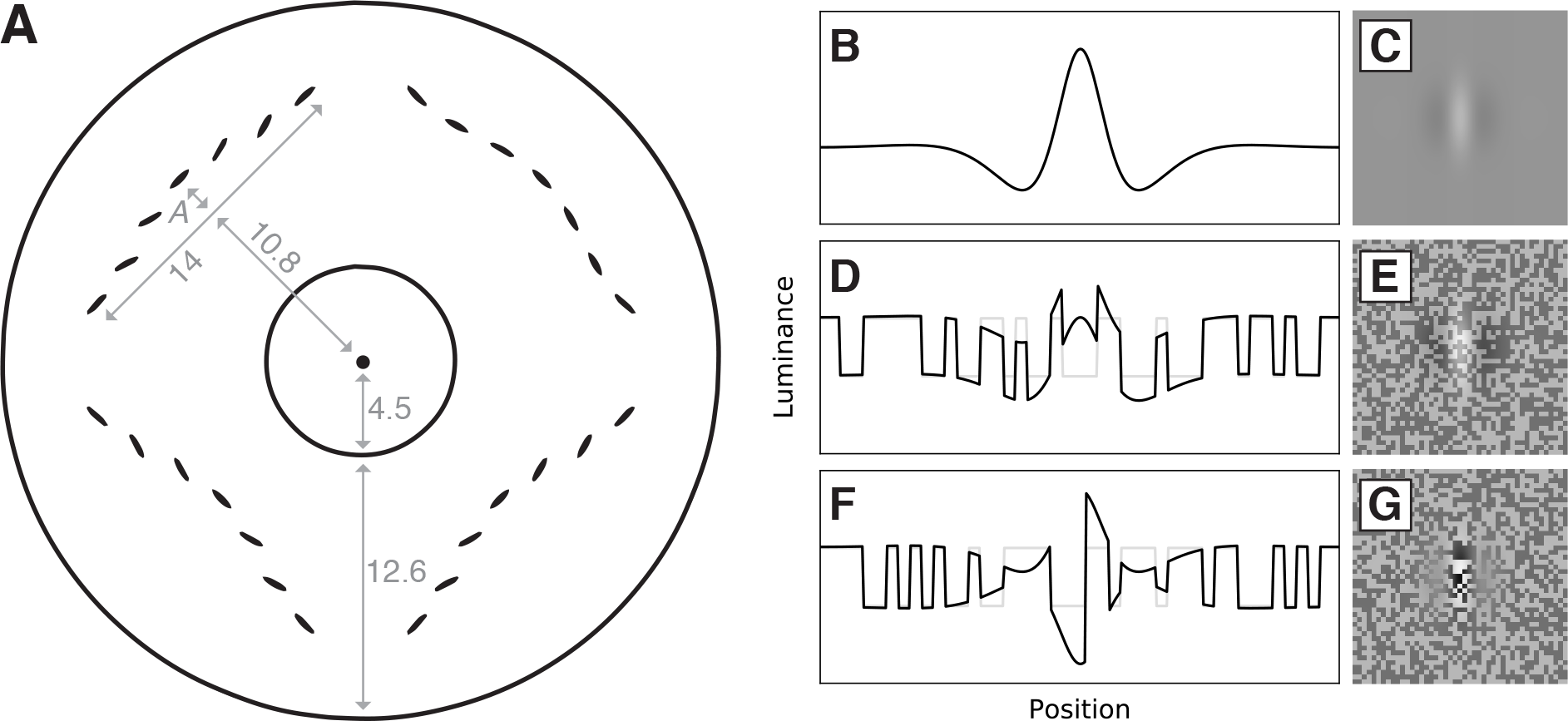
A: Diagram of stimulus layout. Dimensions are given in wavelengths of the wavelets used to form the contours. To give dimensions in degrees of visual angle, these numbers must be divided by the spatial frequency (1.8 c/deg). The central black dot indicates the location of the fixation point. The black circles indicate the onset and offset of the noise texture ring. B: Luminance cross-section of 1st order luminance-modulated log-Gabor wavelet (C). D: Luminance cross-section of 1st order luminance-modulated wavelet added to background of binary noise (E), with the luminance of the noise alone given in light grey. F: Luminance cross-section of 2nd order wavelet (G) formed modulation of binary noise (light grey).

The second-order stimuli require the wavelets to be presented in some background texture. It was possible that the background texture alone could mask the contour stimuli. To account for this, we also measured performance for first-order contours presented on that texture background (Figure 2D-E). The texture was a donut ring shape of binary noise, presented at 30% contrast (Figure 1B-C). The noise was created by scaling up the size of the pixels to form 2×2 pixel blocks. This mitigated the effects of the monitor’s adjacent pixel nonlinearity (Klein et al., 1996). The noise texture ring had its edge blurred by a raised-cosine function, which declined from full- to half-magnitude over 0.9 cycles. Taking the edges as the points where it crosses half-magnitude, the width of the ring was 12.6 cycles.

Second-order contours were generated by multiplying first-order stimuli by the binary white noise ring (Figure 2F-G). Where the first-order wavelets were lighter than mean grey, the second-order wavelets increased the contrast of the texture. Where the first-order wavelets were darker than mean grey, the second-order wavelets reduced the contrast of the texture. The contrast modulation of the second-order wavelets was defined by a contrast-modulation equivalent of the luminance-modulation delta-contrast function (Eq. 2)

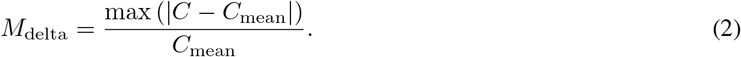

The contrast-modulation is calculated relative to the 30% contrast of our noise texture background. This means that a contrast modulation of 100% would give a stimulus that had a peak contrast of 60% (double the background contrast). The profile of our log-Gabor wavelets has a pronounced peak. That peak is balanced out by a pair of smaller adjacent depressions such that the overall deviation is zero (Figure 2B). We require more headroom above the mean than we do below the mean when we modulate (either luminance or contrast) with our log-Gabor wavelets. Our 30% noise texture contrast therefore allows us to show stimuli with a contrast modulation over 100%.

### 2.4 Procedures

To allow us to equate the visibility of the three types of contours tested in this study (first-order, first-order on noise texture background, second-order modulation of noise texture background) we first measured thresholds for detecting those contours. Each subject’s thresholds were measured using a spatial four-alternative forced-choice task. With the subject fixating centrally, a contour would appear in one of four quadrants (top-right, bottom-right, bottom-left, top-left). The stimulus duration was 800 milliseconds. The subjects had to indicate which quadrant contained the contour. To measure detection thresholds for the first-order contours, their contrast was varied by an adaptive procedure. We used a pair of interleaved staircase routines (Baldwin, 2019). One staircase followed a two-down-one-up rule, and the other followed a one-down-one-up rule. These staircases converged at 71% correct and 50% correct respectively. Following each trial, visual feedback was given to the subject. This indicated whether their response was correct or incorrect (the fixation point either flashed white to indicate “correct”, or black to indicate “incorrect”). Testing was completed when both staircases had reached either 40 trials or 12 reversals. Performance for detecting second-order contours was measured in a similar manner. In this case, the staircases controlled the second-order modulation amplitude. Each subject performed two repetitions of each condition. Thresholds were obtained by fitting the data with a cumulative normal function in Palamedes (Prins and Kingdom, 2009). We combined the data over both repetitions before fitting. These thresholds are presented in Table 1 with 95% confidence intervals obtained using parametric bootstrapping. We used the contrast and modulation thresholds from Table 1 to equate the visibility of the contours in our main experiment. For each subject, the amplitude of the wavelets (either first-order contrast, or second-order contrast modulation) was set to three times their detection threshold. The goal of this was to factor out any difference in simple “visibility” for the different types of modulation.

In the main experiment, we measured the ability of our subjects to discriminate contours with “good continuation”. We defined contours with “good continuation” as those where the orientations and positions of the wavelets forming the contour agreed with the same cosine-shaped curve. This is in contrast to the “distractor” contours, where the orientations of the wavelets were consistent with a contour curving in the opposite direction. The contours in each of the four quadrants could curve either inward (toward fixation) or outward (away from fixation). The direction was chosen randomly for each contour during stimulus generation. Judging which contour had good continuation required the subject to combine the orientation and position information from the wavelets. It would not be possible to discriminate the target from the distractors using only one type of stimulus information.

We again employed a four-alternative forced-choice paradigm with an 800 ms stimulus duration. In each trial, contours were displayed in all four quadrants around the central fixation point. There was one target contour (with good continuation) and three distractors. Following stimulus presentation, subjects made their response by pressing a key on a keypad. They were asked to indicate the quadrant which they believed contained the target stimulus. No emphasis was placed on the speed of response. Subject responses were followed by visual feedback to indicate whether they were correct or incorrect. The difficulty of the task was controlled by the curvature of the contours. In the classic Field et al. (1993) task, curvier contours were more difficult to integrate. However in our task it is *straighter* contours which make the task more difficult. This is because the discrepancy between the position and orientation information in the distractors decreases as the contour becomes straighter. The curvature modulation amplitude was controlled by a pair of staircases (two-down-one-up rule and one-down-one-up). Each staircase was set to terminate after 12 reversals or 50 trials (whichever came first).

We measured noise-masking functions for each wavelet type. These established how curvature modulation thresholds varied when external noise was applied to the contours. For each subject and stimulus condition we set out to measure three repetitions with no external noise. We also measured thresholds with four levels of orientation noise, and with four different levels of position noise. We set out to measure at least two repetitions for each noise level. We chose the external noise standard deviations based on the previous study that established this paradigm (Baldwin et al., 2017). They had found the equivalent internal noise levels for orientation to be 6°. We therefore chose 3°, 6°, 12°, and 17° as the orientation noise standard deviations to test in this study. During stimulus generation, the orientation of each wavelet was randomly jittered. To each wavelet’s orientation, we added an independent sample drawn from a normal distribution. That distribution had a mean of zero and a standard deviation equal to the external noise level being tested. For position noise, Baldwin et al. (2017) found an equivalent internal noise of 3 arcmin (0.3 cycles of their 6 c/deg wavelets). Assuming equivalent internal position noise were to scale with spatial frequency, we would expect our 1.8 c/deg contours in this study to result in an equivalent internal noise of 10 arcmin. We therefore chose values of 7.5, 15, 30, and 42 arcmin for our external position noise standard deviations. Similar to the orientation case, we drew random samples for each wavelet. Two independent samples jittered (separately) the *x* and *y* coordinates of their position.

### 2.5 Curvier contours control experiment

One disadvantage of the contour discrimination task we used in our main experiment (and presented previously in Baldwin et al., 2017) is that the contours tend to be relatively straight around threshold. This would prevent us from drawing conclusions about the discrimination of more curved contours. This can be overcome however, by applying a baseline curvature to the contours (Figure 3). We can consider our basic task to be one where the baseline curvature is zero. The target contour (with “good continuation”) and the distractor contours (where the wavelet orientations and positions are mismatched) are created by shifting the orientations and positions of the wavelets relative to a straight line. In this control experiment, we instead start with a curved baseline contour (randomly curving either inwards or outwards). We then generate our stimuli by shifting the orientations and positions relative to that baseline. For our target contours, we can either shift them to be “less curvy” than the baseline (Figure 3A-B) or to be “more curvy” than the baseline (Figure 3C-D). Crucially, the same sets of orientation and position information are also present in our distractor contours (Figure 3G-J). This prevents subjects discriminating a distractor from a target without combining the orientation and position information together. Achieving this required the “less curvy” and “more curvy” target conditions to be interleaved together. The data from the two conditions were separated for the analysis though, as we expected thresholds may differ between them. We measured thresholds at two baseline curvatures: 1.25 and 1.75 cycles. Three of our four subjects (S1, S2, and S4) took part in this experiment.

**Figure 3:**
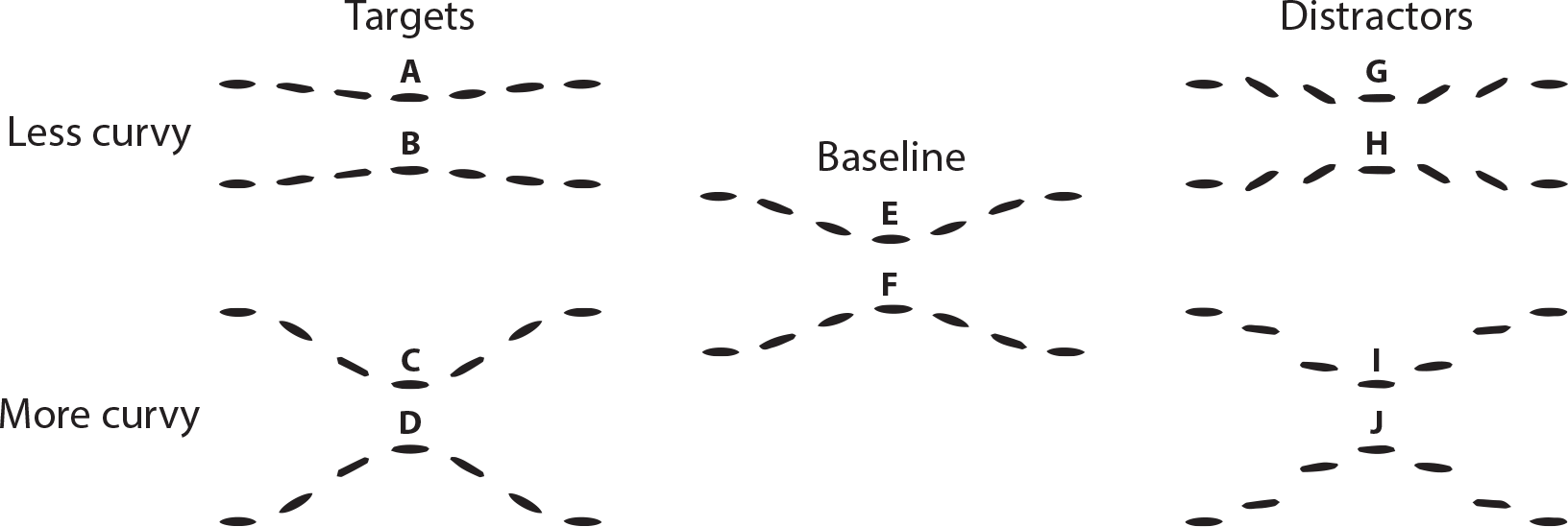
Demonstrating the stimulus design for the control experiment where a baseline curvature was applied to the contours. The scale of the curvatures in this diagram corresponds to a baseline amplitude of 1.75 cycles, with a target modulation amplitude of 1.2 cycles (approximately twice the discrimination threshold for this baseline curvature). The targets are generated by starting with the “baseline” contour (either E or F) and then either shifting the orientations and positions of the wavelets in a way consistent with a less curvy contour (A-B) or consistent with a more curvy contour (C-D). The distractors (G-J) are generated by applying orientation shifts which would be consistent with a change in the *opposite* direction to the position shifts.

### 2.6 Orientation discrimination control experiment

In this study, we present wavelets at three times their contrast or modulation detection thresholds. This was an attempt to equate their visibility. Our method differed from the previous study on second-order contour integration. They instead used an orientation discrimination task to balance the intensity of their first- and second-order wavelets (Hess et al., 2000). We performed a control experiment to determine whether the normalisation method we used in this study also equalises performance in an orientation-discrimination task. Three of our four subjects (S1, S2, and S4) were available to take part in this experiment. The stimulus arrangement is demonstrated in Figure 4. These hexagonal arrangements of seven wavelets could appear at any of the four locations where contours appeared in the main experiment. The centre-to-centre spacing between the wavelets was two cycles. The wavelets are identical to those used in the main experiment. The circular patch of noise was 9 cycles in diameter at half-magnitude. The same softening was applied to its edges as was used in the stimuli in the main experiment. The target was surrounded by a two-pixel thick black ring that was 12.6 cycles in diameter. As in the other experiments, the stimulus duration was 800 ms.

**Figure 4:**
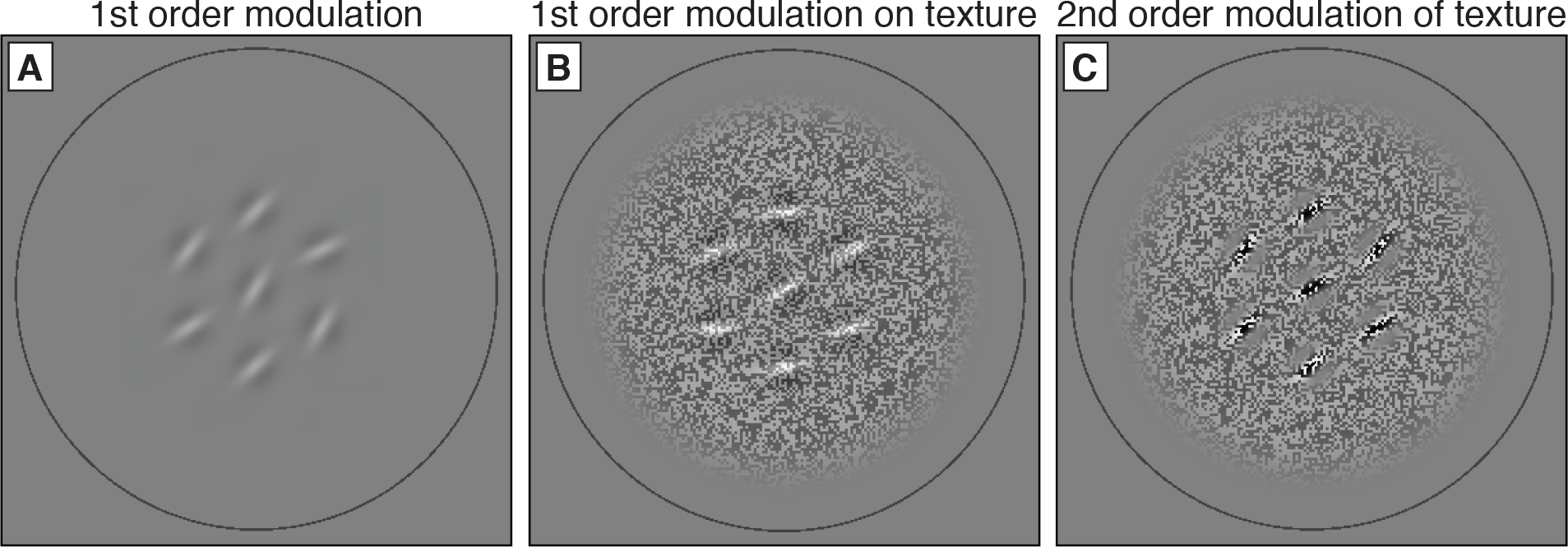
Examples of stimuli presented in the orientation-discrimination control experiment. Panel A shows a 1st order stimulus (at 30% contrast). Panel B shows a 1st order contour (65% contrast) added on top of white noise (30% contrast) . Panel C shows group of 2nd order wavelets that modulate the contrast of that white noise (260% modulation). Each stimulus was rendered with 10° orientation noise standard deviation.

Contrary to the other experiments, thresholds were measured using a temporal two-interval forced-choice design. Before each trial, the black ring would appear at one of the four potential target locations. This cued the subject to attend to that location, while keeping their fixation in the centre of the display. The subject would then confirm they were ready with a button press. Following that confirmation, two stimuli would be shown one after the other (with a blank 200 ms inter-stimulus interval between them). One stimulus (the non-target) would have an orientation between 1 and 135°. The target stimulus would have an additional clockwise rotation (controlled by a staircase). These two stimuli were shown in a random order. The subject’s task was to identify the interval in which the target was presented (where the stimulus was rotated more clockwise). There is an alternate and equivalent description of this task. It can instead be considered as asking the subject to discriminate the orientation of the second (test) stimulus. On each trial, they must determine if it is rotated clockwise or counter-clockwise relative to the first (reference) stimulus. There were four staircases used to control the magnitude of the target rotation. Each staircase controlled the target rotation for one location where the stimuli could be presented. For the condition where the wavelets were defined by first-order modulation added to the texture background, we only measured thresholds without any external noise. For the first-order condition without the background and for the second-order condition we also measured noise-masking functions. For this we made measurements with 7°, 10°, and 15° of external orientation noise. The noise was added to the wavelets in the same way as in the main contour task.

### 2.7 Analysis

For our contour integration task, we obtained thresholds by fitting psychometric functions using Palamedes (Prins and Kingdom, 2009). We used the cumulative normal as the shape of our psychometric functions. These functions define the probability of correctly identifying the quadrant containing the contour with good continuation, as a function of the curvature modulation amplitude. The guess rate was 25% for this four-alternative forced choice task. We assumed a 1% lapse rate. We calculated thresholds at the 55.2% correct point of the psychometric function. This is the percent-correct where *d*’ = 1. In our analysis, we first fit thresholds from individual repetitions of the experiment separately. We calculated the standard error of the threshold using parametric bootstrapping (1,000 samples). Occasionally, there would be a large standard error associated with the threshold. In these cases the subject’s performance did not adequately constrain the function fit. We set a rule to remove repetitions where the standard error of the log_2_ threshold was greater than 1, and to re-run those conditions. The proportion of rejected repetitions ranged from 0% (subject S2) to 16% (subject S4). The data from the repetitions that met the inclusion criterion (standard error less than 1) were merged together. These merged data were then fit to calculate the threshold used in the further analysis. The number of repetitions included for each condition was almost always at least two. The only exception was subject S1 in the first-order stimulus condition with the highest level of position noise. In this case alone, only a single repetition was collected with acceptable data.

We plotted thresholds as a function of external noise standard deviation. This gives the noise-masking function. We fitted the linear amplifier model (Lu and Dosher, 2008)

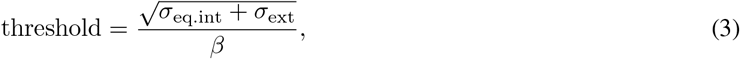

where *σ*_ext_ is the external noise standard deviation, *σ*_eq.int_ is the fitted equivalent internal noise, and *β* is the processing efficiency. The equivalent internal noise *σ*_eq.int_ gives the point at which the noise-masking function transitions from flat to increasing. The efficiency *β* controls the vertical offset of the function. To obtain estimates of efficiency relative to the ideal observer, we used predictions generated from the model presented in our previous publication (Baldwin et al., 2017). We were then able to normalise the *β* values obtained in the current experiment by dividing them by the ideal observer *β*. We obtained the error associated with our LAM parameter estimates through bootstrapping. We re-fitted the model curves to the set of 1,000 bootstrapped sample thresholds generated in the psychometric function fitting. The standard deviation of the bootstrapped parameter values gives the within-subject standard errors. These are reported in Tables 2–3. The standard errors of the between-subject means were calculated by dividing the standard deviation of the log_2_-transformed individual subject values by the square root of the number of subjects. The number of repetitions varied between different subjects and conditions. The minimum number of measurements (total number of repetitions for all noise levels) included for single noise-masking function was 10, the maximum was 20, and the average (both mean and median) was 14. For the second experiment (with the baseline curvature applied to the contours), we did not exclude any data. As the “more curvy” and “less curvy” target conditions were interleaved within each block we obtained only half as much data per repetition (compared to the main experiment). For this reason we did not fit the data from the individual repetitions. We collected between 3 and 5 repetitions for each psychometric function.

For the orientation discrimination control experiment, the psychometric functions were fit in a similar manner to that used in the contour integration experiments. As this was a two-alternative forced-choice task however, the guess rate was 50% and thresholds were calculated at 76% (for *d*’ = 1). We tested whether performance was equal at the four potential stimulus locations using the PAL_PFLR_ModelComparison function from the Palamedes toolbox (Prins and Kingdom, 2009). For each subject and stimulus condition, we compared two approaches to modelling the noiseless threshold data. In the lesser model, the data from all four locations was fit by the same psychometric function. The fuller model allowed there to be different thresholds for stimuli at the different locations. For subject S1 there was never sufficient evidence to reject the lesser model (with significance level *α* = 0.05). For S2, the fuller model was significantly better than the lesser model for the second-order condition (transformed likelihood ratio = 9.0, *p* = 0.034). The threshold fit from the lesser model was 6°. With the fuller model, the thresholds ranged from 4° in the top right to 10° in the top-left. For S3, the fuller model was significantly better than the lesser model for the first-order condition on the noise texture background (transformed likelihood ratio = 16.1, *p* = 0.001). The threshold fit from the lesser model was 9°. With the fuller model, the thresholds ranged from 6° in the bottom right to 21° in the top-left. Of these two significant results, only the latter would be likely to survive appropriate correction for multiple comparisons. As we are not (in this study) interested in variations in sensitivity over the visual field, we combine data across the four locations for the analysis presented in the results section below.

**Table 2:** Fitted linear amplifier model (Eq. 3) parameters for the orientation noise condition. The log_2_-transformed parameter values for each subject (S1-S4) and their mean are given with their standard error. For the individual subject data the standard error is calculated from the bootstrapped parameter estimates. The bottom row gives the mean value converted back to linear units.

| Subject | Internal noise $\log_2(\sigma_{\text{int}})$ | | | Efficiency $\log_2(\beta/\beta_{\text{ideal}})$ | | |
| --- | --- | --- | --- | --- | --- | --- |
| | 1<sup>st</sup> | 1<sup>st</sup> + texture | 2<sup>nd</sup> × texture | 1<sup>st</sup> | 1<sup>st</sup> + texture | 2<sup>nd</sup> × texture |
| S1 | $3.0 \pm 0.4$ | $2.3 \pm 0.3$ | $3.6 \pm 0.3$ | $-0.9 \pm 0.3$ | $-1.5 \pm 0.2$ | $-1.9 \pm 0.2$ |
| S2 | $2.7 \pm 0.2$ | $2.7 \pm 0.2$ | $3.5 \pm 0.2$ | $-1.1 \pm 0.1$ | $-1.4 \pm 0.1$ | $-1.9 \pm 0.2$ |
| S3 | $2.8 \pm 0.2$ | $2.9 \pm 0.2$ | $4.0 \pm 0.6$ | $-1.1 \pm 0.1$ | $-1.2 \pm 0.2$ | $-1.5 \pm 0.5$ |
| S4 | $2.5 \pm 0.2$ | $3.2 \pm 0.2$ | $3.8 \pm 0.5$ | $-1.4 \pm 0.1$ | $-1.4 \pm 0.1$ | $-2.6 \pm 0.4$ |
| Mean | $2.8 \pm 0.1$ | $2.8 \pm 0.2$ | $3.7 \pm 0.1$ | $-1.1 \pm 0.1$ | $-1.4 \pm 0.1$ | $-2.0 \pm 0.2$ |
|  | 7° | 7° | 13° | 46% | 39% | 25% |

The noise-masking functions for the orientation discrimination control task were fit in a similar manner to the main contour task. We applied the same rejection rule for standard errors greater than 1. The proportion of rejected repetitions ranged from 0% to 13%. The number of repetitions used for a single masking function ranged from 6 to 9 (mean and median were both 8). We mathematically derived the ideal observer prediction for our orientation discrimination control task (see Appendix 6). We use this to plot the the ideal observer prediction in Figure 8, and normalise the subject’s efficiencies to be relative to the ideal observer in Figure 9 and Table 6.

## 3 Results

### 3.1 Discrimination of good continuation

Figure 5 shows the noise-masking functions. Curvature-modulation thresholds are averaged over four subjects. The two rows show results obtained with our two different external noise types. The top row shows the effect of adding orientation noise to the wavelets. The bottom row shows the effect of adding position noise. The three columns show results from three stimulus types: first-order contours on a mean-grey background, first-order contours on a binary noise texture background, and second-order contours generated by modulating the contrast of the binary noise texture. These correspond to the three contour images shown in Figure 1. The left-most point in each subplot gives the threshold measured when no external noise was added to the contours. For this reason, those points are duplicated across the two rows of subplots. The difference between thresholds measured for first-order contours with and without the noise texture background was small. For second-order contours however the threshold was more than a factor of two higher. It is worth noting that we *are* able to measure thresholds for second-order contours. This distinguishes our findings from those of Hess et al. (2000), who found little measurable ability to integrate second-order contours. That study used the Field et al. (1993) task where the subject must detect smooth contours hidden in a background of random wavelets.

**Figure 5:**
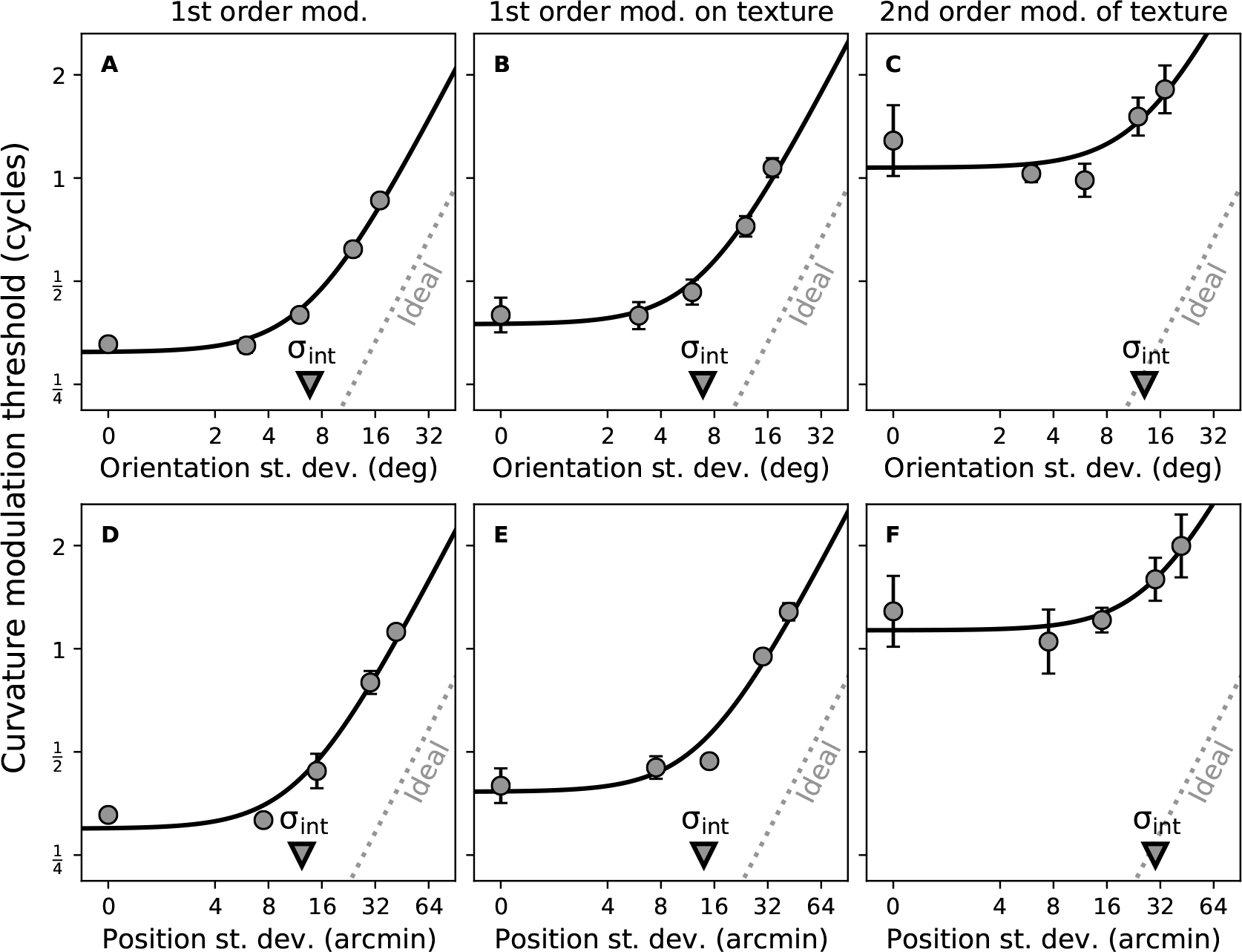
Curvature modulation thresholds required to discriminate the smooth contour, plotted against the external orientation or position noise added to the wavelets composing the contour. Thresholds are averaged across four subjects, with error bars showing standard error. Data are fitted with the linear amplifier model (Eq. 3). The fitted equivalent internal noise parameter is indicated by the downward-pointing triangle symbol on the x-axis. The diagonal grey line indicates the performance of the ideal observer model Baldwin et al. (2017).

When the variance of the external noise is small, thresholds do not appear to be affected. This results in a flat noise-masking function between the first two points on each subplot in Figure 5. Beyond some critical point however, increasing the level of external noise causes a proportional increase in threshold. Thresholds for first-order contours on the binary noise background are consistently a little higher than those without the background. For second-order contours the difference is more dramatic, with a large increase in threshold (although the difference is reduced at the higher external noise variances). We performed a two-way ANOVA (in R; R Core Team, 2014; RStudio Team, 2016) on the thresholds presented in each row of Figure 5. The two factors were the contour type (first-order, first-order on texture, or second-order modulation of texture) and external noise standard deviation. For the orientation noise condition, we found significant effects of contour type (*F*_2,6_ = 61.7, *p* < 0.001) and external noise standard deviation (*F*_3,9_ = 143.1, *p* < 0.001). We also found a significant interaction between the two (*F*_6,18_ = 3.5, *p* = 0.018). For the position noise condition, we also found significant effects of contour type (*F*_2,6_ = 22.8, *p* = 0.002) and external noise standard deviation (*F*_3,9_ = 88.1, *p* < 0.001). We again found a significant interaction between the two (*F*_6,18_ = 6.6, *p* < 0.001). We can therefore state that adding external noise to our stimuli significantly increases thresholds. This gives us a noise-masking function for each of the three noise types. Additionally, these three noise-masking functions significantly differ from each other. For further analysis we apply a model that describes the shape of these functions.

The noise-masking functions in Figure 5 are fit by the Linear Amplifier Model (Equation 3). The black curves are determined by the two fitted parameters of that model. The equivalent internal noise (*σ*_int_) determines the point at which the function transitions from flat to increasing. Its value is marked with a triangle symbol on the *x*-axis of each subplot. In both rows, the equivalent internal noise approximately doubles between the first-order and second-order subplots. The second fitted parameter is the processing efficiency (*β*). This determines the location of the linear part of the masking function. A benchmark value is given by the ideal observer model. This indicates the performance expected on the task from a model which makes best possible use of the available information. An ideal observer model for this task was defined previously by Baldwin et al. (2017). The ideal observer model is assumed to perfectly recover the positions and orientations of all of the wavelets in the display. Its behaviour is indicated by the diagonal dashed grey lines in each subplot. The performance of our human subjects relative to the ideal observer can be judged by the distance between those lines and the increasing part of the fitted black curves. Performance is closer to the ideal observer for the first-order conditions compared to the second-order conditions. As the shape of the fitted curves is determined entirely by the two LAM parameters (equivalent internal noise and processing efficiency), we conduct our main analysis on the values of those fitted parameters.

The fitted model parameters averaged over our four subjects are presented in Figure 6. The top and bottom rows again give the data from the orientation noise and position noise conditions respectively. The two columns give the fitted equivalent internal noise (*σ*_int_) and processing efficiencies (*β*). The processing efficiency is calculated relative to the ideal observer. This means that 100% is the theoretical best possible performance on the task. As seen from the triangle markers in Figure 5, the equivalent internal noise values for the two first-order conditions in Figure 6A and Figure 6C are similar. We find an equivalent internal orientation noise of 7° for our first-order stimuli, similar to the 6° value reported in the previous study that used this task (Baldwin et al., 2017). For the equivalent internal position noise, the value we find here is higher (12 arcmin, compared to 3 arcmin in the previous study). Expressed relative to the stimulus spatial frequency however, the two become similar (0.30 vs. 0.36 cycles).

**Figure 6:**
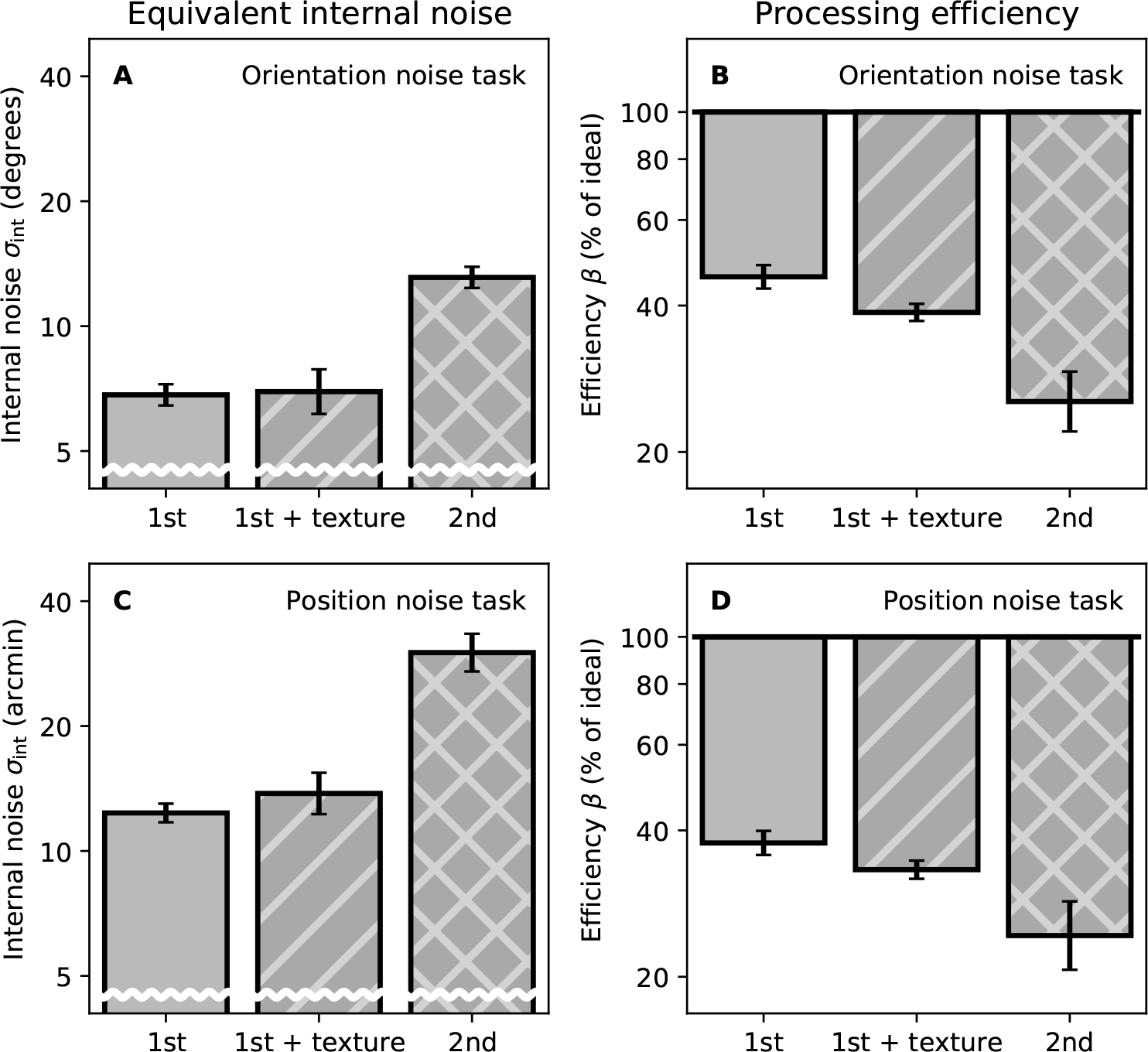
Parameters obtained by fitting the linear amplifier model (Eq. 3) to the data from the four individual subjects. The average parameters (geometric mean) across subjects are presented with their standard errors.

For both the orientation and position noise-masking functions, the equivalent internal noise for the second-order condition is roughly twice that of the two first-order conditions. We investigated this with a pair of one-way ANOVAs. For each noise type, we looked at the log-transformed equivalent internal noise across the different stimulus conditions (using the individual subject data shown in the left half of Tables 2 and 3). We found significant effects for both our orientation noise (*F*_2,6_ = 12.0, *p* = 0.008) and our position noise (*F*_2,6_ = 25.9, *p* = 0.001) masking functions. In the pairwise comparisons (uncorrected paired t-tests), the differences between the two first-order conditions were non-significant in both cases (*p* = 0.937 for the orientation noise condition, *p* = 0.574 for position). The ANOVA effect was driven by a significant increase in second-order equivalent internal noise (*p* = 0.008 comparing first-order to second-order for orientation, *p* = 0.009 for position). We can therefore state that at least part of the reduced sensitivity to second-order contours is due to factors leading to an increased equivalent internal noise.

**Table 3:**
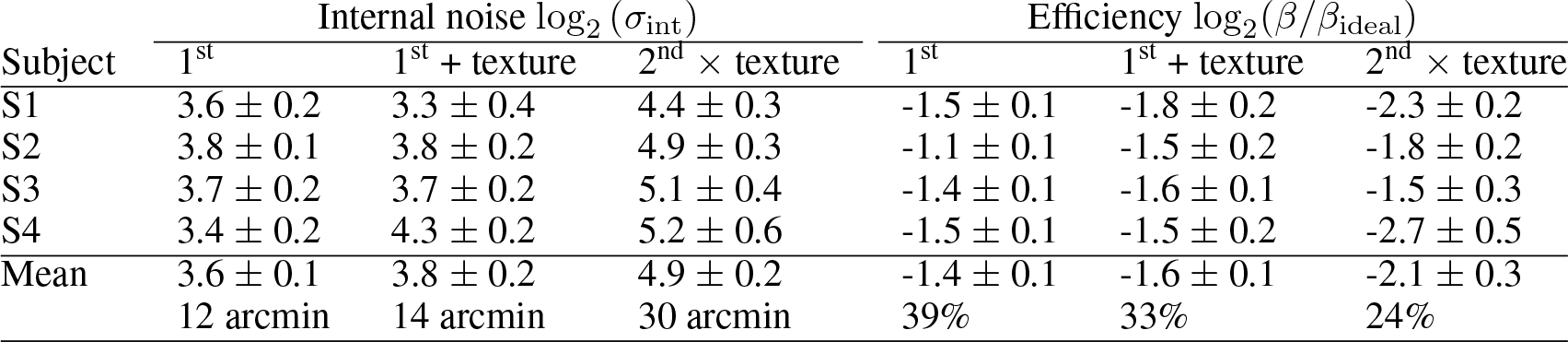
Fitted linear amplifier model parameters for the position noise condition. For further details see Table 2 caption.

| Subject | Internal noise $\log_2(\sigma_{\text{int}})$ | | | Efficiency $\log_2(\beta/\beta_{\text{ideal}})$ | | |
| --- | --- | --- | --- | --- | --- | --- |
| | 1<sup>st</sup> | 1<sup>st</sup> + texture | 2<sup>nd</sup> × texture | 1<sup>st</sup> | 1<sup>st</sup> + texture | 2<sup>nd</sup> × texture |
| S1 | $3.6 \pm 0.2$ | $3.3 \pm 0.4$ | $4.4 \pm 0.3$ | $-1.5 \pm 0.1$ | $-1.8 \pm 0.2$ | $-2.3 \pm 0.2$ |
| S2 | $3.8 \pm 0.1$ | $3.8 \pm 0.2$ | $4.9 \pm 0.3$ | $-1.1 \pm 0.1$ | $-1.5 \pm 0.2$ | $-1.8 \pm 0.2$ |
| S3 | $3.7 \pm 0.2$ | $3.7 \pm 0.2$ | $5.1 \pm 0.4$ | $-1.4 \pm 0.1$ | $-1.6 \pm 0.1$ | $-1.5 \pm 0.3$ |
| S4 | $3.4 \pm 0.2$ | $4.3 \pm 0.2$ | $5.2 \pm 0.6$ | $-1.5 \pm 0.1$ | $-1.5 \pm 0.2$ | $-2.7 \pm 0.5$ |
| Mean | $3.6 \pm 0.1$ | $3.8 \pm 0.2$ | $4.9 \pm 0.2$ | $-1.4 \pm 0.1$ | $-1.6 \pm 0.1$ | $-2.1 \pm 0.3$ |
|  | 12 arcmin | 14 arcmin | 30 arcmin | 39% | 33% | 24% |

Comparing efficiency across the three stimulus conditions, the pattern is again similar between the orientation noise (Figure 6B) and the position noise (Figure 6D) masking results. For the first-order stimulus condition, efficiency was higher for the orientation noise condition (46%) compared to position noise condition (39%). This was also true in the previous study (Baldwin et al., 2017), although they found a larger difference (36% vs. 21%). Additionally, the efficiencies measured in this study are higher overall compared to those measured previously. This may be due to our larger stimulus scale. For both the orientation and position noise conditions, there appears to be a slight decrease in efficiency when the first-order stimuli are placed on the binary noise texture background. Further decreases are seen for the second-order stimulus condition. We performed one-way ANOVAs using the data from the right half of Tables 2 and 3. We compared fitted processing efficiency parameters across the different stimulus conditions. We found a significant effect for the orientation noise case (*F*_2,6_ = 12.2, *p* = 0.008). There was no significant pairwise comparison between the two first-order conditions (*p* = 0.152). There was however a significant difference between first- and second-order (*p* = 0.021). For the position noise case there was not a significant effect of stimulus condition (*F*_2,6_ = 4.6, *p* = 0.062). Therefore, we can also attribute part of the reduced sensitivity to second-order contours as being due to a reduced efficiency with which the orientation information of the wavelets is processed. The results with position noise do trend in the same direction, but do not achieve statistical significance.

### 3.2 Extension of task to more curved contours

Results from the task with the baseline curvature applied to the contours are shown in Figure 7. Results from the first experiment are included (baseline curvature of zero) for reference. Thresholds for second-order stimuli remain higher than those for first-order stimuli. A quick comparison can be made across each of the four pairs of data points measured in this experiment. Thresholds are consistently around three times higher for second-order contours. Surprisingly, thresholds are consistently lower for more curvy targets than for less curvy targets. This is true both for first-order and second-order contours. For first-order contours there is a tendency for thresholds to increase with baseline curvature, though this is more apparent with the “less curvy” targets. For second-order contours, any such relationship is less clear.

**Figure 7:**
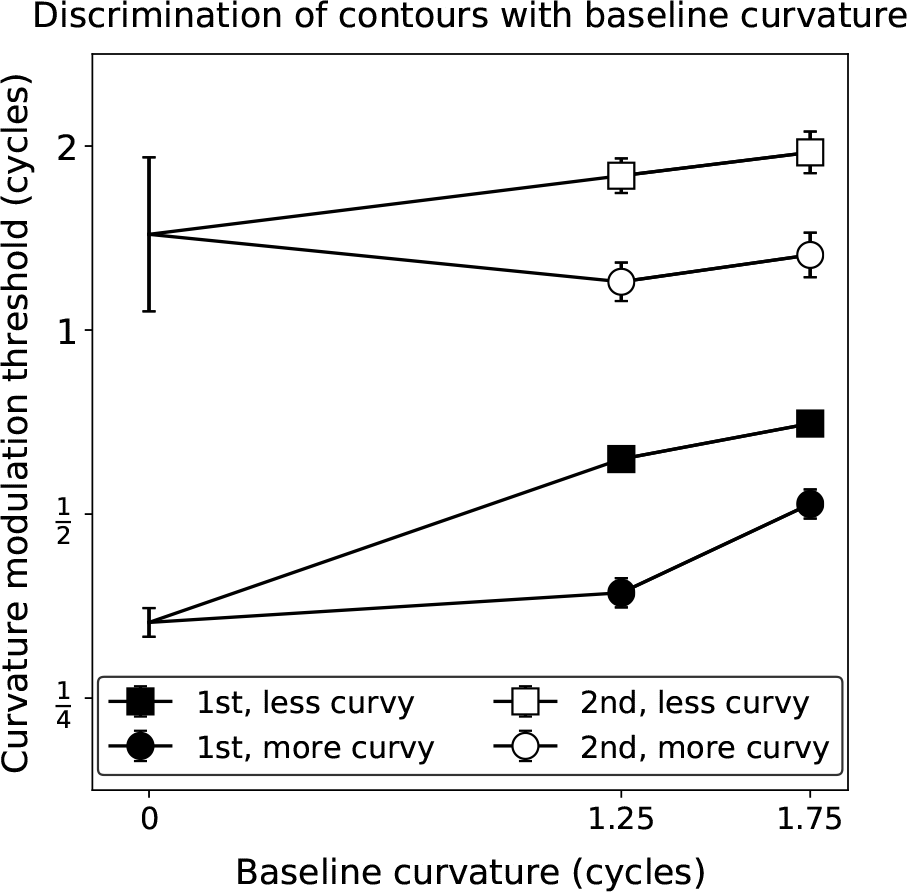
Results from the experiment where a baseline curvature was applied to the contours. Thresholds presented are averaged over three subjects (S1, S2 & S4), with error bars giving the standard error. Thresholds with zero baseline curvature are taken from the main experiment data.

**Figure 8:**
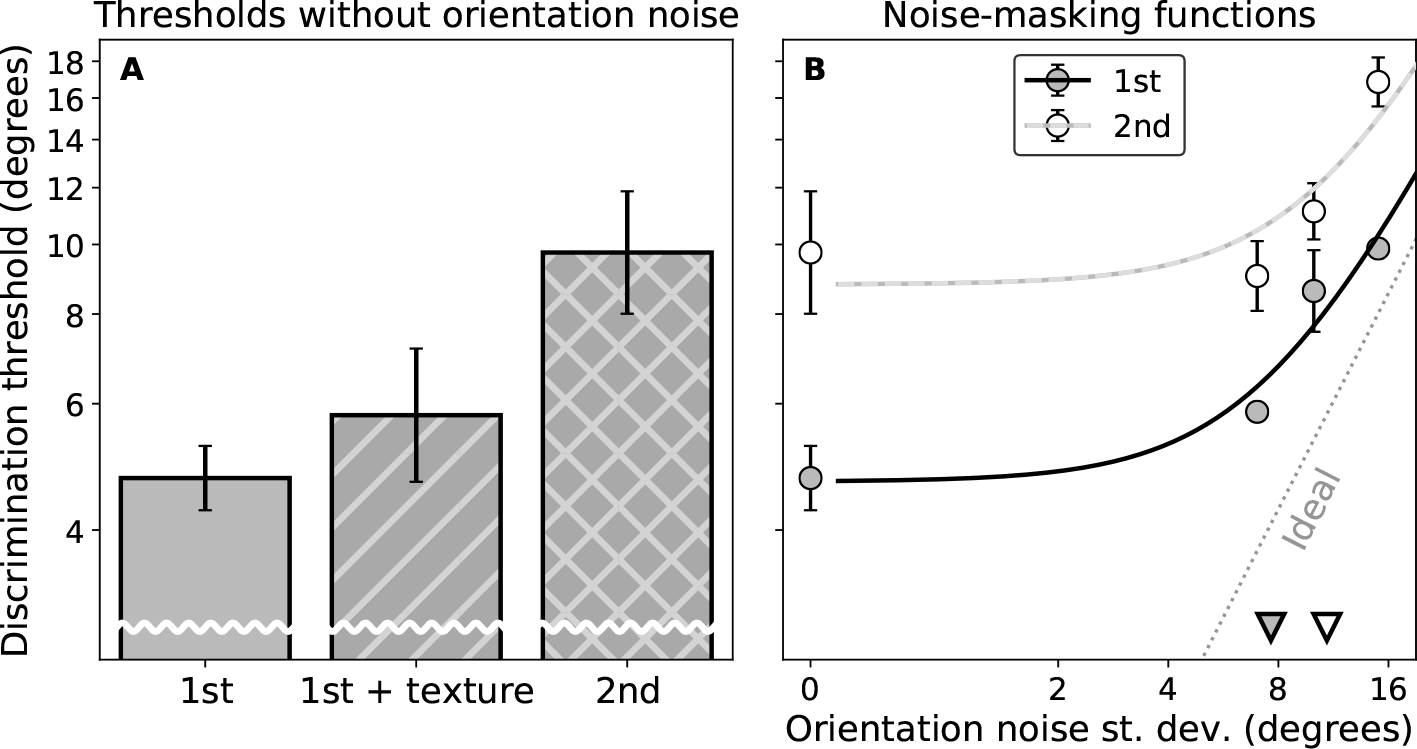
Results from the orientation discrimination control experiment. Panel A shows the thresholds obtained without any external noise added to the stimuli. Panel B shows the noise-masking functions measured for the 1^st^ order and 2^nd^ order stimuli. The dashed diagonal grey line gives the expected performance of the ideal observer. The solid lines give the fits of the linear amplifier model (Eq. 3) to the data. The triangle markers indicate the equivalent internal noise parameter values from those fits.

**Figure 9:**
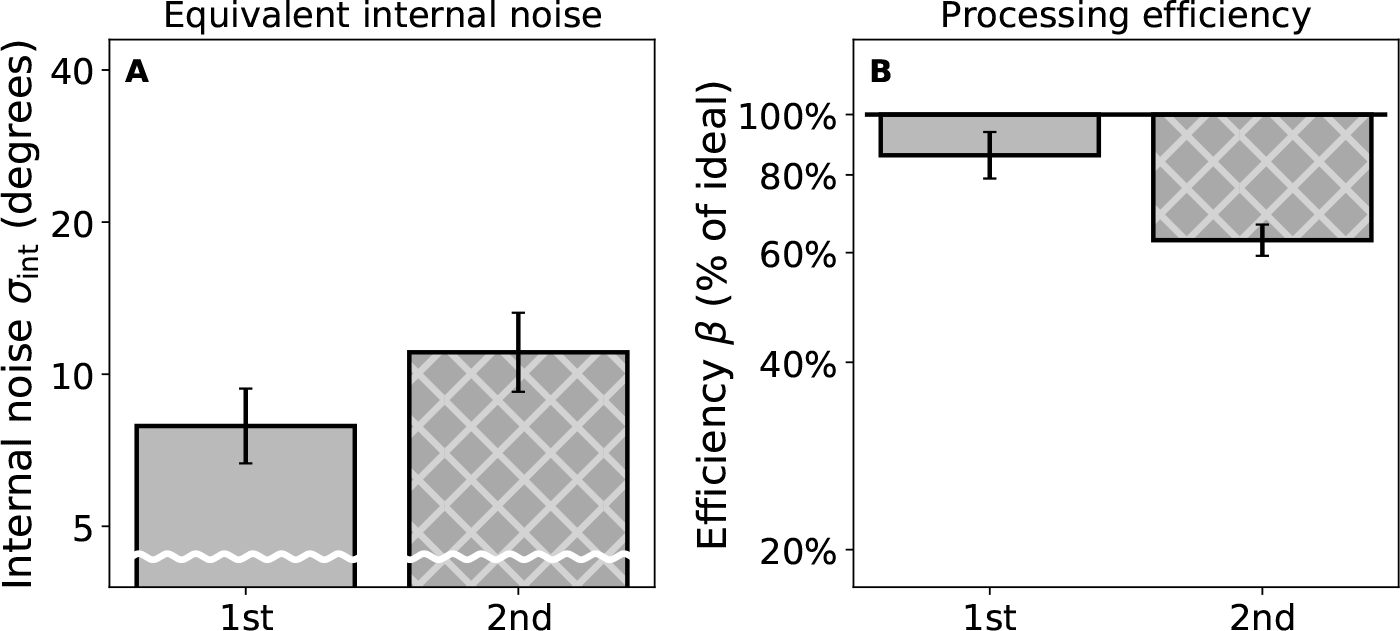
Parameters obtained by fitting the linear amplifier model (Eq. 3) to the data from the orientation discrimination control experiment. Data are averaged over three individual subjects (geometric mean). Error bars give standard errors.

Results from individual subjects are given in Tables 4 and 5. It is worth noting that, in the second-order condition, average thresholds for the “less curvy” targets are greater than the magnitude of the baseline curvature. Because in this condition the contours are defined as a *reduction* in curvature from that baseline, that means that the targets at threshold here are actually slightly curved in the opposite direction (having passed through being a straight line, where the target curvature is equal to the baseline). A three-way ANOVA was performed on the thresholds. The factors were the baseline curvature (1.25 or 1.75 cycles), the stimulus condition (first- or second-order), and whether the target contour was less or more curvy than the baseline. We found significant effect of stimulus condition (*F*_1,2_ = 150.1, *p* = 0.007) and curvature direction (*F*_1,2_ = 158.7, *p* = 0.006). There was no significant effect of baseline curvature (*F*_1,2_ = 7.6, *p* = 0.110). There were also no significant interactions. We conclude that second-order contours can be integrated even when they are curved. The performance disadvantage relative to first-order contours remains. Surprisingly, we find a performance *advantage* (both in the first- and second-order results) for detecting more curved contours. This is contrary to the results from a previous study (using the paradigm from Field et al., 1993), which found the integration of more curved contours to be impaired in the periphery (Hess and Dakin, 1999).

**Table 4:** Curvature modulation thresholds measured in the experiment where a 1.25 cycle baseline curvature was applied to the contours. Results are given from the three individual subjects (log_2_ thresholds), along with the mean (plotted in Figure 7). The value of the mean converted back to linear units is also given.

| Subject | log<sub>2</sub> thresholds for 1.25 cycle baseline |  |  |  |
| --- | --- | --- | --- | --- |
|  | Less curvy |  | More curvy |  |
|  | 1<sup>st</sup> order | 2<sup>nd</sup> order | 1<sup>st</sup> order | 2<sup>nd</sup> order |
| S1 | -0.81 ± 0.09 | 1.04 ± 0.09 | -1.53 ± 0.36 | 0.51 ± 0.21 |
| S2 | -0.64 ± 0.08 | 0.64 ± 0.07 | -1.23 ± 0.19 | 0.19 ± 0.19 |
| S4 | -0.66 ± 0.08 | 0.84 ± 0.08 | -1.52 ± 0.23 | 0.08 ± 0.30 |
| Mean | -0.70 ± 0.05 | 0.84 ± 0.12 | -1.43 ± 0.10 | 0.26 ± 0.13 |
|  | 0.6 cycles | 1.8 cycles | 0.4 cycles | 1.2 cycles |

**Table 5:** Curvature modulation thresholds measured in the experiment where a 1.75 cycle baseline curvature was applied to the contours. For further details see Table 4 caption.

| Subject | $\log_2$ thresholds for 1.75 cycle baseline | | | |
| --- | --- | --- | --- | --- |
|  | Less curvy |  | More curvy |  |
|  | 1<sup>st</sup> order | 2<sup>nd</sup> order | 1<sup>st</sup> order | 2<sup>nd</sup> order |
| S1 | $-0.42 \pm 0.08$ | $1.13 \pm 0.07$ | $-0.75 \pm 0.09$ | $0.54 \pm 0.30$ |
| S2 | $-0.57 \pm 0.11$ | $0.69 \pm 0.14$ | $-1.02 \pm 0.14$ | $0.11 \pm 0.20$ |
| S4 | $-0.54 \pm 0.11$ | $1.08 \pm 0.08$ | $-1.07 \pm 0.23$ | $0.57 \pm 0.28$ |
| Mean | $-0.51 \pm 0.05$ | $0.97 \pm 0.14$ | $-0.95 \pm 0.10$ | $0.41 \pm 0.15$ |
|  | 0.7 cycles | 2.0 cycles | 0.5 cycles | 1.3 cycles |

**Table 6:** Fitted equivalent internal noise and processing efficiency parameters for the orientation discrimination control experiment. Parameters were obtained by fitting the linear amplifier model (Eq. 3). Results are given for the three individual subjects, along with their average.

| Subject # | Internal noise $\log_2(\sigma_{\text{int}})$ | | Efficiency $\log_2(\beta/\beta_{\text{ideal}})$ | |
| --- | --- | --- | --- | --- |
|  | 1<sup>st</sup> order | 2<sup>nd</sup> order | 1<sup>st</sup> order | 2<sup>nd</sup> order |
| S1 | $2.6 \pm 0.3$ | $4.0 \pm 0.7$ | $-0.5 \pm 0.2$ | $-0.5 \pm 0.7$ |
| S2 | $2.7 \pm 0.2$ | $2.9 \pm 0.3$ | $-0.3 \pm 0.2$ | $-0.6 \pm 0.2$ |
| S4 | $3.6 \pm 0.4$ | $3.5 \pm 0.5$ | $0.1 \pm 0.3$ | $-0.9 \pm 0.4$ |
| Mean | $3.0 \pm 0.3$ | $3.5 \pm 0.3$ | $-0.2 \pm 0.2$ | $-0.7 \pm 0.1$ |
| | $8^\circ$ | $11^\circ$ | 85% | 63% |

### 3.3 Comparison with orientation discrimination

Thresholds obtained in the orientation discrimination control experiment are shown in Figure 8A. A one-way ANOVA found a significant effect of stimulus condition (*F*_2,4_ = 9.2, *p* = 0.032). There were no significant pairwise comparisons however (comparing the two first-order conditions gives *p* = 0.320, comparing first-against second-order gives *p* = 0.073). As the first-order results with and without the background noise texture were similar, we only measured noise masking functions for the first- and second-order stimulus conditions. These are shown in Figure 8B. A two-way ANOVA was performed. It found significant effects of both stimulus condition (*F*_1,2_ = 23.3, *p* = 0.040) and external noise level (*F*_3,6_ = 13.3, *p* = 0.005). There was not a significant interaction between the two (*F*_3,6_ = 2.5, *p* = 0.154). This means that we do have a significant masking effect from our external orientation noise. There is also a significant difference between the results from our first- and second-order stimulus conditions. Our attempt to equate our stimuli by presenting them at multiples of their contrast- or modulation-detection thresholds did not result in equalised performance on an orientation-discrimination task.

The average fitted linear amplifier model parameters for the orientation discrimination control are given in Figure 9. It appears that the relative lack of sensitivity in the second-order task is due to a combination of two factors. There is a small increase in internal noise between the first- and second-order stimulus conditions. There is also a moderate decrease in efficiency. The parameter values from individual subjects reported in Table 6 complicate this story however. Of the three subjects, each shows a different pattern of results. For subject S1 there is a 2.6-fold increase in internal noise between first- and second-order, but no change in efficiency. For subject S2 both parameters show a slight deficit for second-order. There is a 1.1-fold increase in internal noise and a 1.2-fold *reduction* in efficiency. For subject S4 the internal noise is similar for first- and second-order, but there is a moderate (1.4-fold) drop in efficiency. The consequence of these individual differences is that neither the equivalent internal noise (*t*_2_ = *−*1.1, *p* = 0.389), nor the efficiency (*t*_2_ = 1.5, *p* = 0.281) show a significant difference between the first- and second-order stimulus conditions.

## 4 Discussion

We find that subjects are able to discriminate the good continuation of contours formed from second-order contrast-modulation of texture. Performance was inferior to that achieved with first-order luminance modulation. This was the case even when first-order contours were presented on a texture background. We measured noise-masking functions for the different contour types. This allowed us to investigate the reason for the performance differences. We found that equivalent internal noise for second-order contour integration was approximately twice that for first-order. This was true both for the orientation noise condition (13° vs. 7°) and the position noise condition (30 arcmin vs. 12 arcmin). This means that the quality of the second-order input into the contour-integrating mechanism is impaired compared to the first-order input. The properties of these inputs could be further explored by experiments that manipulate the bandwidths of the wavelets forming the contours (Baldwin et al., 2015). It is informative to compare the equivalent internal noise parameters from the current study to those obtained in Baldwin et al. (2017). That study found a similar equivalent internal noise for orientation (6°). For their position noise condition, the equivalent internal noise was much lower (3 arcmin). This can be explained by the difference in spatial frequency between that study (6 c/deg) and the current study (1.8 c/deg). Converting the position noise into cycles (periods of the stimulus spatial frequency) brings the two values into agreement. The equivalent internal noise in Baldwin et al. (2017) was 0.30 cycles, whereas in the current study it was 0.36 cycles (for first-order stimuli). This demonstrates scale-invariance across a three-fold range of spatial frequency. Previous work using the Field et al. (1993) task has also found contour integration to be scale-invariant (Hess and Dakin, 1997).

Our methods allow us to compare results from our contour task against those from our orientation discrimination task. We find similar equivalent internal noise in the two tasks. This differs from the similar comparison made in our previous study (Baldwin et al., 2017). In that study, the equivalent internal noise for contour integration was larger (6° vs. 2°). Importantly, the control condition in this study has subjects judge the orientation of a texture composed of seven wavelets. The control condition in Baldwin et al. (2017) features only a single element. In that previous study, we hypothesised that contour integration may involve a “squelching” of orientation noise. This could result in a thresholding of deviant orientations. Similar phenomena have been seen in studies of orientation variance discrimination (Morgan et al., 2008; Christensen et al., 2015). At the time we noted that there appeared to be no such threshold affecting the position noise data reported in Baldwin et al. (2017). A subsequent noise variance discrimination study conducted using shapes (sampled and re-synthesized with wavelets) also found that the thresholding reported for orientation noise (Christensen et al., 2015) was not present for position noise (Christensen et al., 2019).

The Baldwin et al. (2017) task was developed to measure contour integration as a test of “good continuation” discrimination. Our method does not require the perfomance-limiting noise background used in the Field et al. (1993) task (Watt et al., 2008). However, the results found in this study raise a question: if the only deficit for second-order contours was increased equivalent internal noise (as described mentioned above), would we expect to see the difference in performance reported by Hess et al. (2000)? Adding orientation noise to contours in the Field et al. (1993) task does not dramatically affect performance. A previous study demonstrated this for foveal presentations of the contour stimulus (Hess and Dakin, 1999). Within their stimulus array (10.4 degrees of visual angle square), subjects could still locate the contour (performance greater than 75%) with more than 10° of orientation jitter added to the wavelets. This level of external noise is actually slightly lower than the equivalent internal noise we found for second-order contours. It should be considered however that we made our measurements with contours presented 6 degrees of visual angle from fixation. If subjects were able to make eye movements in our study (as was allowed in Hess et al., 2000) to foveate the contours we would expect the equivalent internal noise to decrease. Therefore the orientation noise should not have impaired performance in Hess et al. (2000). For position noise the Hess and Dakin (1999) study finds performance remains above 75% until at least 0.5 cycles of positional jitter is applied to the target contour. This is lower than our second-order equivalent internal position noise. We may also expect however, that a lower value would have been obtained if subjects viewed the stimuli freely. It is possible that the effects of the equivalent internal orientation and position noise would combine together to affect performance. This may be sufficient to explain the lack of second-order contour integration ability in Hess et al. (2000). A potential future study could adapt our paradigm to look at how the two factors interact (similar to the work performed by Christensen et al., 2019).

We also find differences in processing efficiency between the stimulus conditions. These differences could attribute some of the reduced second-order performance to higher-level factors. We note however that the difference in efficiency between first- and second-order was only statistically significant for the orientation noise condition. It is possible that contour-processing may be less efficient in the Hess et al. (2000) task. They used the Field et al. (1993) method, where contours are presented in a background of random wavelets. Experiments with contours hidden in background noise have found that those contours are extracted by a recurrent processing network (Altmann et al., 2003; Kourtzi et al., 2003; Chen et al., 2014; Drewes et al., 2016; Kuai et al., 2017; Li et al., 2019). This involves feedback from higher cortical areas beyond primary visual cortex. Whether a similar pathway is necessary for the task we present here depends on the role of those recurrent connections. If their purpose is to segregate contours from the background noise (Chen et al., 2014) then they will not be necessary for our task. Our results suggest a feed-forward contour-building process is capable of second-order contour integration. In that case it may be that the feed-back contour-segregation process is deficient. If so, that would explain the difference between our results and those presented in Hess et al. (2000). This accords with results from a recent fMRI study. They found distinct patterns of activation dependent on whether contours were presented within a noisy background context (Qiu et al., 2016).

We normalised the visibility of the different types of contours in this study. We did so by presenting them at three times their detection threshold. For our second-order contours it would not have been possible to show contours of much higher visibility. Our normalisation already rendered them at close to the maximum possible contrast-modulation. So, what is the role of second-order contour integration in normal visual experience? Previous results suggest that first- and second-order vision are multiplexed (Li et al., 2014). If so, then the ability to integrate second-order contours may be due to an activation of mechanisms that are more usefully driven by first-order inputs. On the other hand, it is likely that second-order contour integration is more useful outside of the artificial conditions established here. The task we set to our subjects puts them at a double-disadvantage compared to natural viewing. Both second-order contrast modulation sensitivity (Hess et al., 2008) and first-order contour integration ability (Nugent et al., 2003) decline with eccentricity. We expect that better performance would be seen in a task where the subject could foveate the target. Also, we have not explored cases where contours are defined by a combination of first- and second-order modulation. This would agree with previous work on texture perception. Second-order information may be used to enhance performance, when compared to the use of first-order information alone (Johnson et al., 2007). Our study re-opens the book on the integration of second-order contours. Further work will be required to explore these nuances.

## 5 Additional information

## 5.1 Acknowledgements

This research was funded by an NSERC Discovery Grant (RGPIN-2016-03740) awarded to Robert F. Hess. This pre-print manuscript was written in LATEX using TeXShop. It is formatted with a custom style available at: github.com/alexsbaldwin/biorxiv-inspired-latex-style.

## 5.2 Author contributions

The study was conceived and designed by ASB, MK and RFH. The experiment was conducted by ASB and MK. The analysis was performed by ASB. The manuscript was drafted by ASB. All authors provided revisions to the manuscript.

## 6 Appendix: Ideal observer model for orientation discrimination control experiment

Here we derive the ideal observer prediction for our orientation discrimination control task. In the first interval, a reference stimulus is shown composed of seven wavelets. The *i*^th^ wavelet has its orientation drawn from a Normal distribution, represented as 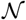 (mean, variance), to give us

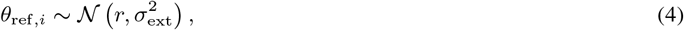

where *r* is the reference orientation, and *σ*_ext_ is the standard deviation of the orientation noise. Technically the orientation distribution is circular, and so the Von Mises distribution would be more appropriate for this modelling. This is unnecessary however, as even the maximum standard deviation we consider (15^°^) is relatively small (only one in a million randomly sampled orientations would be > 75° from the reference orientation). We then show the test stimulus in the second interval. Each *i*^th^ wavelet has the orientation

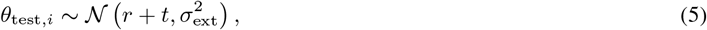

where *t* is the expected orientation difference. The subject must discriminate the direction of this difference (positive or negative). The ideal observer solution to this task is the one which chooses the response with the greatest likelihood (Green and Swets, 1966; Geisler, 2004). The maximum likelihood estimators of *r* and *r* + *t* are the sample means in the reference and test intervals. We can therefore achieve ideal performance on this task by comparing the mean orientations from the test 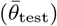 and reference 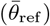 intervals. If 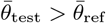 then the test stimulus is rotated clockwise relative to the reference (and so the expected orientation difference *t* is most likely positive). We can calculate the difference in the average orientations as

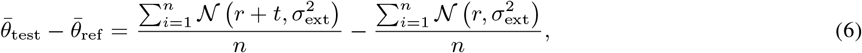

where *n* = 7 for our seven samples, this simplifies to

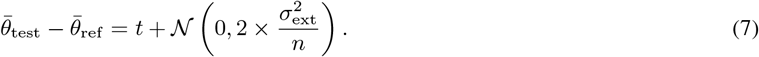

We can use this to calculate the ideal observer’s signal to noise ratio (*d*’ = *μ/σ*) for this task

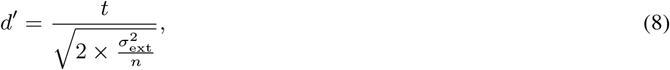

solving for *d*’ = 1 we have

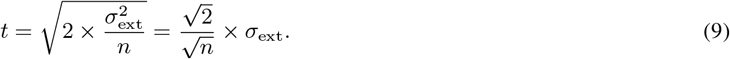

By substituting in to Equation 3, we can also calculate the ideal observer’s *β* in the LAM (their *σ*_eq.int_ is zero by definition) which is 1.87, or (as we compare log-transformed parameters in our tables) we can take the log_2_ of that, which gives us 0.90.

